# ComCat: Combating Covariate Effects in Brain Analysis

**DOI:** 10.64898/2026.06.18.733200

**Authors:** Christian Gaser, Robert Dahnke, Habib Ganjgahi, Thomas E. Nichols

## Abstract

As neuroimaging analysis shifts toward large-scale, multi-site studies, managing the unwanted variability introduced by combining heterogeneous datasets has become a critical challenge. Although tools such as ComBat and its neuroimaging extensions are widely used to address this variability, they only permit the modeling of categorical site effects and cannot account for continuous sources of confounding, such as image quality, head motion, and acquisition parameters. We introduce ComCat, an extension of the ComBat framework that preserves biologically relevant covariates while removing the effects of categorical site indicators and continuous nuisance variables. The latter are modeled as smooth nonlinear functions via B-spline basis expansion. ComCat is applicable to a broad range of brain analysis tasks, including voxel- and surface-based morphometry, normative modeling, and machine learning–based prediction. To demonstrate its capabilities, we evaluated ComCat on brain age prediction across five datasets covering complementary multi-site harmonization scenarios: ON-Harmony (10 subjects × 6 scanners; n = 80); the Buchert traveling-phantom dataset (1 subject × 116 scanners; n = 531); the Tohoku single-scanner, varying-acquisition dataset (n = 121); MR-ART (148 subjects with varying motion levels); and an ABIDE subset comprising 229 control subjects and 208 individuals with autism spectrum disorder across 14 scanners. Using image quality measures derived from CAT12 as continuous nuisance variables, ComCat reduced the mean absolute error (MAE) in brain age prediction relative to ComBat-GAM in all five datasets, including the two scenarios where site information was unavailable or uninformative. In the ABIDE dataset, ComCat improved harmonization while preserving the difference between the control and ASD groups, demonstrating that scanner-related variance can be removed without affecting biologically meaningful signals. ComCat can operate with or without site labels and is agnostic to the source of image quality metrics.

## Introduction

Multi-site neuroimaging studies are essential for achieving the large sample sizes and demographic diversity required by modern brain analysis methods. However, these studies introduce scanner-related variability that can obscure biological signals. This variability stems from variations in field strength, pulse sequences, receiver coil characteristics, and software versions among imaging sites (Fortin et al., 2017; Fortin et al., 2018) and impacts the entire spectrum of downstream analyses, including voxel- and surface-based morphometry (VBM and SBM), regional volumetric analyses, normative modeling, and machine learning–based prediction frameworks, such as brain age (Cole and Franke, 2017; Franke et al., 2010; Franke and Gaser, 2019; Marquand et al., 2016; Rutherford et al., 2022). Without harmonization, comparisons across sites can produce spurious effects that mimic disease-related changes or mask genuine biological differences (Lombardi et al., 2020; Pomponio et al., 2020).

ComBat (Johnson et al., 2007) is the most widely used method of harmonization in neuroimaging. Originally developed for correcting batch effects in microarray gene expression data, ComBat has been adapted for diffusion tensor imaging (Fortin et al., 2017) and cortical thickness measurements (Fortin et al., 2018). It has since been applied across a broad range of neuroimaging modalities. ComBat models additive and multiplicative site effects using an empirical Bayes framework that borrows information across features to stabilize parameter estimates. This makes ComBat robust even for small per-site sample sizes. Several extensions have been proposed, including ComBat-GAM, which accounts for nonlinear effects of preserved covariates (Pomponio et al., 2020); LongComBat, which handles longitudinal data (Beer et al., 2020); CovBat, which harmonizes cross-feature covariances (Chen et al., 2022); and ComBatLS, which preserves covariate effects on both location and scale (Gardner et al., 2024).

However, variability related to the scanner or acquisition process cannot be fully captured by a discrete site label. Variation within sites arising from head motion, susceptibility artifacts, incomplete fat suppression, suboptimal shimming, or differences in spatial resolution introduces additional confounding factors that are continuous in nature and cannot be modeled as categorical batch effects. Growing evidence suggests that image quality systematically affects brain morphometry estimates and, consequently, brain age predictions. Lower image quality is associated with less grey matter and might reflect in elevated brain age (BA) values (Reuter et al., 2015; Nárai et al., 2024; Gilmore et al., 2021). This confounding factor is particularly problematic when image quality correlates with clinical status. For example, patients with neuropsychiatric disorders tend to move more during scanning, which results in lower-quality images that may be misinterpreted as an increase in grey matter loss or greater apparent aging. Most directly, Moqadam et al. (2024) recently showed that head motion during scanning systematically inflates predicted brain age, with the magnitude of the bias scaling proportionally with motion severity, so that motion-corrupted scans appear neurobiologically older than they are.

The standard ComBat method can only correct for discrete site effects and cannot account for continuous confounding parameters. To overcome this limitation, we created ComCat, which is an expansion of the ComBat harmonization framework. ComCat simultaneously preserves biologically relevant covariates (e.g., age and group) while eliminating the impact of categorical site indicators and continuous nuisance variables, including image quality measures (IQMs). To capture nonlinear relationships between IQMs and imaging features, ComCat models each nuisance variable as a B-spline basis expansion within a generalized additive model (GAM). This concept is analogous to ComBat-GAM (Pomponio et al., 2020), but it solves a different problem. While ComBat-GAM uses smooth nonlinear modeling for preserved covariates (e.g., age), ComCat applies it to nuisance covariates that are sources of unwanted variation.

ComCat is applicable to many neuroimaging analyses that require harmonization as a preprocessing step. These analyses include voxel- and surface-based morphometry, regional volumetric analyses, and machine learning–based prediction frameworks. To demonstrate its performance and properties, we evaluate ComCat’s performance and properties on brain age prediction using five datasets with complementary multi-site designs: a controlled multi-subject traveling-phantom dataset (ON-Harmony), an extreme single-subject multi-scanner dataset (Buchert), a single-scanner varying-acquisition dataset (Tohoku), a dataset with varying levels of motion artifacts (MR-ART), and a heterogeneous clinical multi-site cohort (ABIDE). Brain age prediction is an informative test case because, in single-subject and traveling-phantom designs, all variation in predicted age reflects technical sources. This provides a direct readout of harmonization effectiveness. In multi-site clinical samples, brain age prediction tests whether scanner variance is removed and whether biologically meaningful group differences are preserved simultaneously.

## Methods

### ComCat: Theoretical framework

Like ComBat, ComCat is based on the idea that each measured imaging feature can be broken down into a biological signal of interest, technical bias terms, and residual error. ComBat estimates one site bias term (a discrete shift and scaling per site) while preserving the desired biological signal. Then, it subtracts the site bias to produce harmonized data. ComCat extends this concept by incorporating a second class of bias terms derived from continuous sources, such as IQMs. Since the relationship between a continuous variable and an imaging feature does not have to be linear, ComCat models each continuous nuisance variable as a smooth function (a B-spline) of that variable, similar to GAMs.

The harmonization itself proceeds in three conceptual steps. First, the data are centered: an estimated grand mean and the contribution of the preserved biological covariates (e.g., age) are subtracted, and the result is divided by an estimate of the pooled standard deviation. This puts every feature on a common scale and isolates the technical bias terms. Second, ComCat estimates the additive site bias (per-site shifts) and the smooth contribution of each continuous nuisance variable on the standardized data and removes both, so the technical bias terms vanish as expected. Per-site variance differences are also corrected at this stage. Third, the data are rescaled to their original units. The previously subtracted biological covariate contribution and grand mean are added back, and the result is multiplied by the pooled standard deviation. Therefore, the output lies in the same numerical range as the input, but with site and image-quality variation removed and the biological signal preserved.

In the next subsection, we formally describe the data model and the parameter estimation procedure.

### Data model

ComCat extends the location/scale model underlying ComBat (Johnson et al., 2007; Fortin et al., 2017) by partitioning the design matrix into three components. For feature *v*(*v*= 1, …, *P*, e.g., voxels or brain regions) and sample *j* ( *j*= 1, …, *N*), the model is:

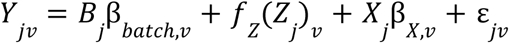

where *B* is the *N* × *K* dummy-coded batch indicator matrix (*K* = number of sites; reduced to a single intercept column when no site information is provided), *Z* is the *N* × *Q* matrix of continuous nuisance covariates (e.g., IQMs), *f*_*Z*_ denotes a smooth nonlinear function of *Z* realized through B-spline basis expansion (see below), and *X* is the *N* × *R* matrix of covariates of interest to be preserved (e.g., age, group). The batch indicators *B* and nuisance covariates *Z* together represent the unwanted bias to be removed, while *X* contains the biological effects to be preserved and ε is the error term.

### B-spline expansion of continuous nuisance variables

Each continuous nuisance variable *z* is expanded into a B-spline basis of dimension *d*:

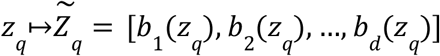

Where *b*_1_, …, *b*_*d*_ are B-spline basis functions defined over equally spaced knots within the observed range of *z*_*q*_ . The basis dimension is the standard GAM smoothness parameter, denoted gam_df_ throughout this paper. To balance flexibility and overfitting risk while adapting to sample size, gam_df_ is selected automatically as:

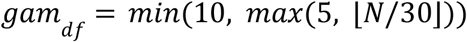

The lower bound of 5 ensures that nonlinear shape can be captured even in small samples; the upper bound of 10 limits flexibility to prevent overfitting. The implications of this upper bound for preserving biological group differences are addressed in the Discussion. The expanded nuisance design matrix is denoted 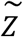with *Q*_‘_ = *Q* · *d* columns.

### Parameter estimation

All parameters are estimated jointly in a single ordinary least squares (OLS) step on the full design matrix:

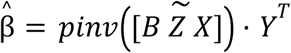

By the Frisch-Waugh-Lovell theorem, the estimate 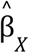 for the preserved covariates is identical whether obtained from this joint model or from a regression on residuals after partialling out 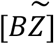. The preserved covariates are therefore correctly accounted for regardless of their correlation with batch or nuisance variables, provided the design matrix is not rank-deficient.

### Standardization

The grand mean is defined as the sample-average predicted value from the batch and nuisance components:

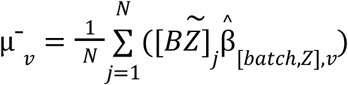

The pooled standard deviation is the root mean squared error of the full-model residuals:

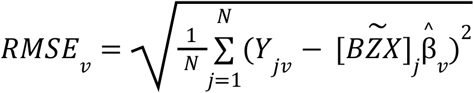

Standardized data are obtained by first removing the grand mean and then the preserved covariate effects. Finally, the data are divided by the pooled standard deviation:

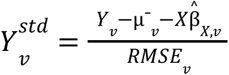

### Estimation of nuisance effects on standardized data

After standardization, the additive effects of the batch and nuisance variables are re-estimated using the reduced design matrix and the standardized data 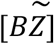:

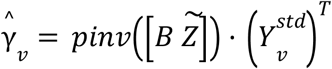

Site-specific multiplicative effects are estimated directly from within-site variances of standardized data:

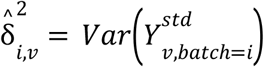

Estimating 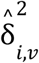 from the within-site variance directly (without first removing the additive nuisance effects) preserves the full per-site variance structure that the multiplicative correction is intended to remove. If the mean-only option is set, 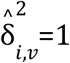 for all batches.

### Data adjustment

For each batch *i*, the adjustment is:

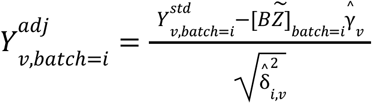

The subtraction term removes both the batch-specific additive shift and the smooth nuisance contributions simultaneously, since the full 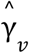 vector is applied via the combined design Matrix 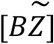.

### Rescaling to original units

The adjusted data are rescaled back to the original data space:

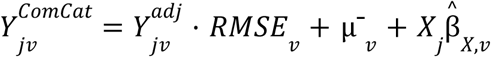

This restores the original data scale while preserving the biological effects encoded in *X*.

### Scale correction: categorical vs. continuous nuisance variables

In ComBat, the multiplicative batch effect δ_*iv*_ captures systematic differences in data variance across sites. For a categorical batch this is well-defined, since all samples within a batch share the same variance scaling. ComCat retains this per-batch scale correction.

The situation is fundamentally different for continuous nuisance variables. Since a continuous variable does not partition the sample into discrete groups, there is no natural analog of within-batch variance. A genuine variance correction along a continuous axis would require modeling the conditional variance, for example, via local variance estimation in *Var* 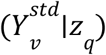 sliding windows or parametric heteroscedasticity models. In neuroimaging applications, however, IQMs primarily affect the mean rather than the variance of derived imaging measures. The added complexity of a heteroscedasticity model, such as bandwidth selection, parametric assumptions, and instability at distribution tails, is not warranted for typical use cases. Therefore, ComCat applies scale correction only for categorical batch effects and limits the treatment of continuous nuisance variables to location correction.

### Relationship to ComBat and its extensions

When no continuous nuisance variables are specified, i.e., when Z is empty, ComCat reduces algebraically to ComBat, but with the Empirical Bayes (EB) step disabled. The omission of EB is a deliberate design choice, not a limitation. Johnson et al. (2007) introduced EB to stabilize batch parameter estimates when per-batch sample sizes are very small. Their examples had as few as four samples per batch, a common situation in early microarray experiments. In neuroimaging, however, per-site sample sizes typically exceed 20 subjects, at which point OLS estimates of γ and δ^2^ are already well-determined and EB shrinkage adds negligible benefit. The EB posterior mean converges to the OLS estimate as batch size grows. Moreover, the EB framework cannot be extended to continuous nuisance variables, which lack the discrete grouping structure required for pooling information across features within a batch.

### Image preprocessing

We processed structural MRI data using CAT12.9 (Gaser et al., 2024). Affine registration of gray matter segmentations was performed using the MNI152Nlin2009cAsym template. We resampled these segmentations to spatial resolutions of 4 and 8 mm, then smoothed them using Gaussian kernels with full widths at half maximum (FWHM) of 4 and 8 mm. This yielded four feature representations per image.

### Image quality measures

Seven IQMs derived from CAT12 (Gaser et al., 2024; Dahnke et al., 2025) were used as continuous nuisance variables in ComCat (Table 1). These IQMs capture complementary aspects of image quality that are known to affect estimates of brain morphometry, even in high quality datasets without visible motion artifacts (Kalc et al., 2026). Although CAT12-derived IQMs were used for compatibility with the preprocessing pipeline, the ComCat framework is agnostic to the source of nuisance variables and can accommodate IQMs from other tools, such as MRIQC (Esteban et al., 2017).

**Table 1.**
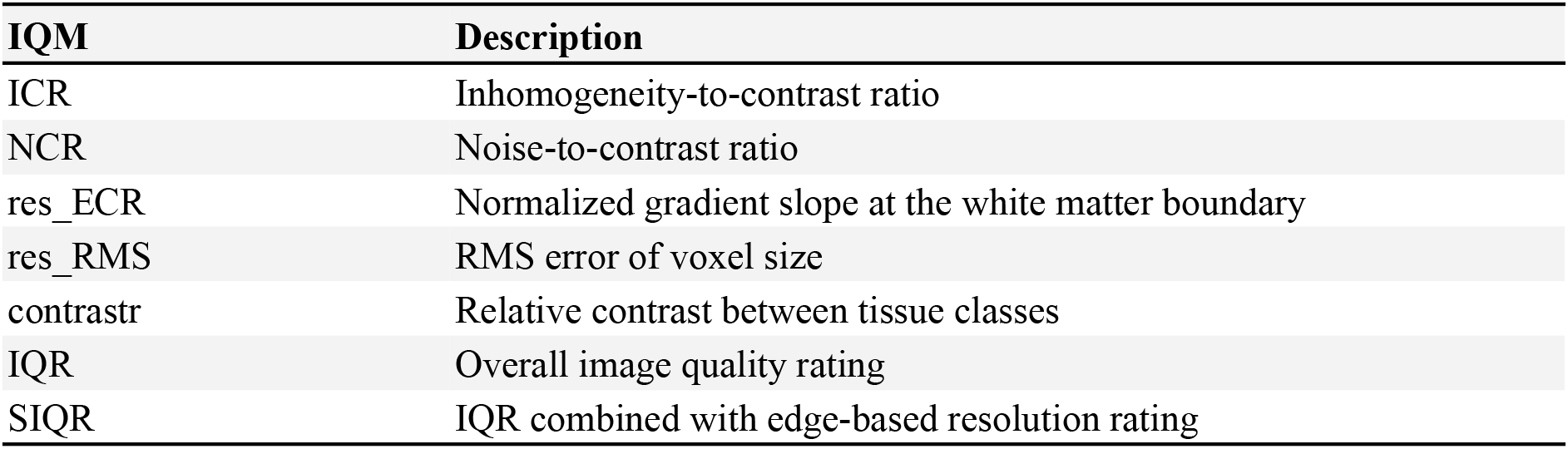
Overview of CAT12 image quality parameters.

### BrainAGE estimation

Brain age was estimated using the BrainAGE framework (Franke & Gaser, 2019; Kalc et al., 2024). Gaussian process regression (GPR) was applied to each of the four feature representations (4 mm/8 mm resampling combined with 4 mm/8 mm smoothing). Predictions were then averaged using weights that were inversely proportional to the squared mean absolute error (MAE). The predictions were age-bias-corrected as described in Smith et al. (2019).

Normative training samples were chosen separately for each evaluation dataset to find a tradeoff between large training sample size and matched age range (Table 2). The data used for the normative training sample for adults is described in Kalc et al. (2024). Source code for BrainAGE is publicly available at https://github.com/ChristianGaser/BrainAGE.

**Table 2.**
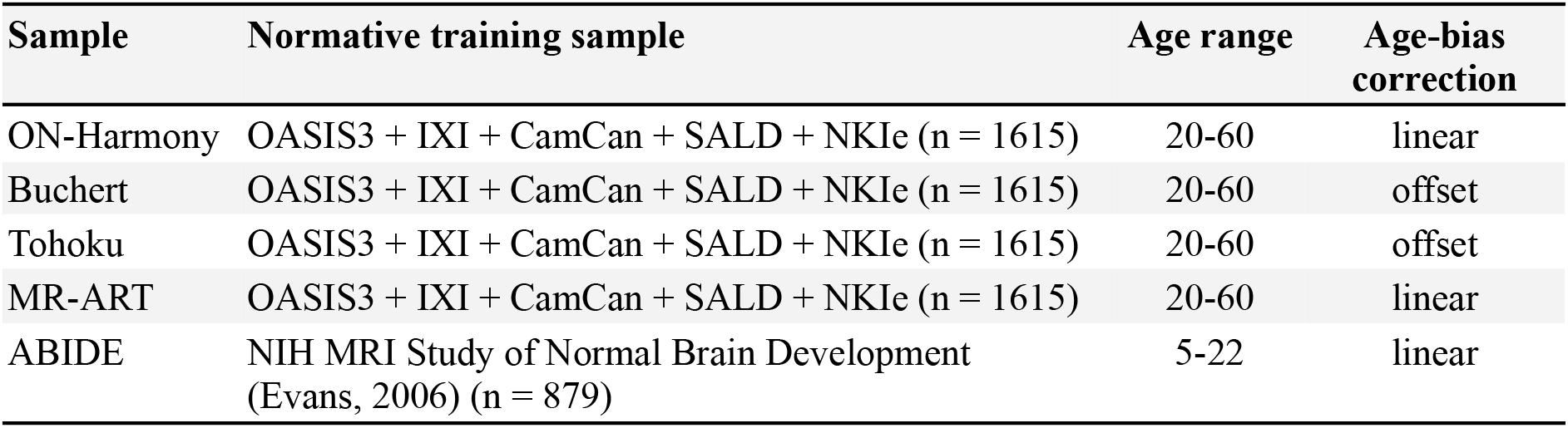
Overview of normative training samples and BrainAGE parameters.

| Sample | Normative training sample | Age range | Age-bias correction |
| --- | --- | --- | --- |
| ON-Harmony | OASIS3 + IXI + CamCan + SALD + NKIe (n = 1615) | 20-60 | linear |
| Buchert | OASIS3 + IXI + CamCan + SALD + NKIe (n = 1615) | 20-60 | offset |
| Tohoku | OASIS3 + IXI + CamCan + SALD + NKIe (n = 1615) | 20-60 | offset |
| MR-ART | OASIS3 + IXI + CamCan + SALD + NKIe (n = 1615) | 20-60 | linear |
| ABIDE | NIH MRI Study of Normal Brain Development (Evans, 2006) (n = 879) | 5-22 | linear |

### Evaluation metric

We use the mean absolute error (MAE) in brain age prediction computed against chronological age as the primary metric for harmonization performance. In single-subject or test-retest designs (e.g., ON-Harmony, Buchert, and Tohoku), where chronological age essentially remains quite similar across scans, all observed variability in predicted brain age reflects technical variation (e.g., scanner, acquisition, and image quality) rather than biological variation. Therefore, a lower MAE in these settings directly indicates that more variance attributable to site, image quality, and motion artifacts has been removed. In multi-site clinical samples (ABIDE), where chronological age varies, a lower MAE in control subjects under preserved age and group covariates indicates that scanner-related variance has been removed without affecting the biological age signal. Thus, MAE offers a concise and interpretable single-number summary of harmonization effectiveness.

### Evaluation samples

Five datasets were selected to span complementary multi-site harmonization scenarios (Table 3). Typical resolutions were about 1 mm, acquired in about five minutes, with sagittal orientation. The TE/TR/TI were around 3/2000/900 ms, with flip angles of approximately 8° and 15° (Table 4).

**Table 3.** Basic overview of evaluation samples.

| Sample | Subjects | Scanners | Scans | Age range | Design |
| --- | --- | --- | --- | --- | --- |
| ON-Harmony | 10 | 6 | 80 | 24-48 | Traveling phantom, multi-subject |
| Buchert | 1 | 116 | 531 | 49-51 | Traveling phantom, single-subject |
| Tohoku | 1 | 1 | 121 | 35 | Single scanner, varying acquisition |
| MR-ART | 148 | 1 | 436 | 18-75 | Three levels of motion artefacts |
| ABIDE | 437 | 14 | 437 | 10-18 | Clinical multi-site (controls + ASD) |

**Table 4.**
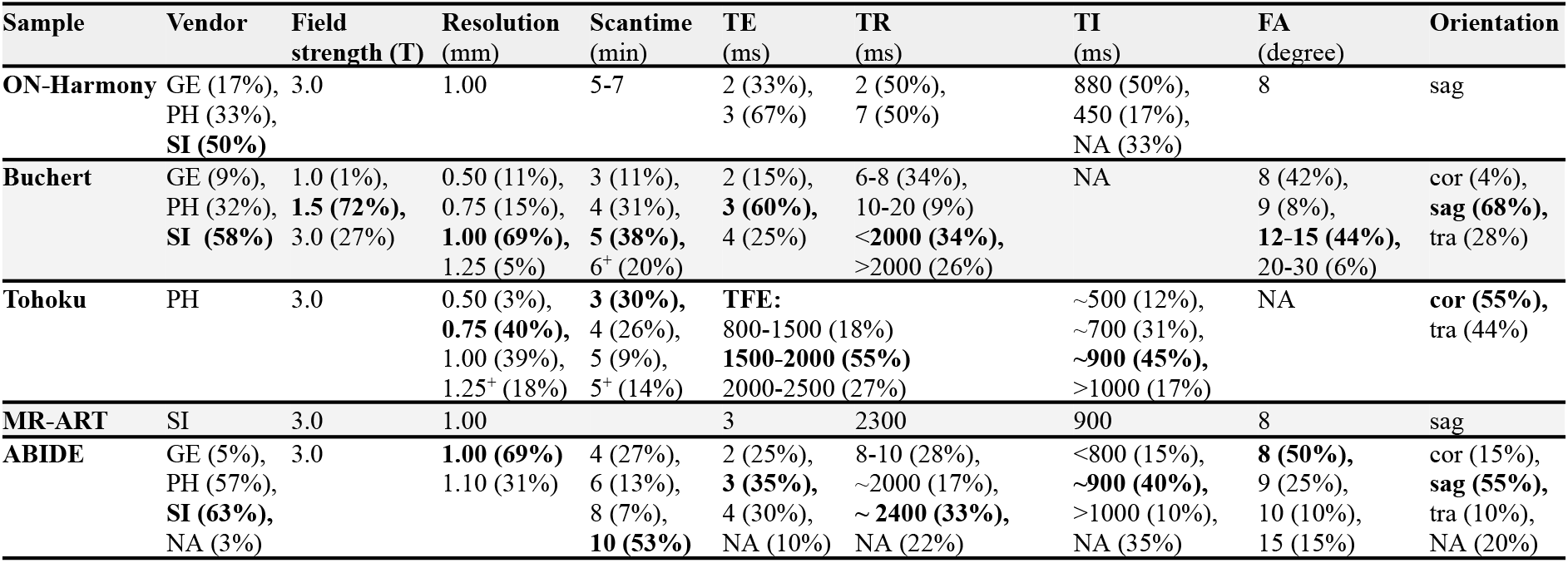
Technical overview of imaging parameter with manufacturer/vendor (GE: General Electrics, PH: Philips, SI: Siemens), field strength, (rounded) average resolution, approximated scantime, echo/relaxation/inversion times (TE/TR/TI), flip angle (FA), and orientation (sag: sagittal, cor: coronal, tra: transversal). NA=not available

**Table 5.** Brain age prediction MAE (years) by harmonization condition. The lowest MAE estimates are displayed in bold.

| Sample | Raw | ComBat-GAM | ComCat (IQMs) | ComCat (IQMs + site) |
| --- | --- | --- | --- | --- |
| ON-Harmony | 4.103 | 3.546 | 1.773 | **1.658** |
| Buchert | 2.334 | 0.904 | 1.019 | **0.698** |
| Tohoku | 5.001 | - | **0.982** | - |
| MR-ART | 3.845 | - | **3.100** | - |
| ABIDE (control subjects) | 1.361 | 1.173 | 0.984 | **0.971** |

#### ON-Harmony

(Hawco et al., 2025, Warrington et al., 2023; https://openneuro.org/datasets/ds004712) is a multi-site, multi-subject traveling-phantom dataset comprising 10 participants scanned 80 times across 6 scanners (ages 24-48). It is the most controlled evaluation scenario, allowing direct attribution of within-subject between-scanner variation to scanner effects.

#### Buchert

(Buchert et al., 2025; https://www.kaggle.com/datasets/ukeppendorf/frequently-traveling-human-phantom-fthp-dataset; Opfer et al., 2023) is an exceptional single-subject traveling-phantom dataset: 1 participant scanned 531 times over 2 years across 116 scanners (ages 49-51), with 3-11 repeated scans per scanner. Most scans were acquired at private radiological practices and small hospitals, providing a wide range of acquisition quality representative of real-world clinical heterogeneity.

#### Tohoku

(Thyreau et al., 2013) consists of 121 scans of a single subject (age 35) acquired with systematically varying parameters (acquisition time 36-611 s, in-plane resolution 0.5-1.9 mm, slice thickness 0.6-2 mm) on a single scanner, isolating the effects of acquisition parameters and image quality without any site or scanner change.

#### MR-ART

(Nárai et al., 2022); https://openneuro.org/datasets/ds004173/versions/1.0.2 consists of 363 scans of 148 subjects (ages 18-75) acquired while they were staying still and while they were moving their heads slightly and excessively (i.e., up to three sets of data for each participant) on a single scanner.

#### ABIDE

(Di Martino et al., 2014; https://fcon_1000.projects.nitrc.org/indi/abide/) is a heterogeneous clinical multi-site dataset. We used a subset of ABIDE-I restricted to ages 10-18 years comprising 229 healthy controls and 208 individuals with autism spectrum disorder (ASD) across 14 scanners (85% male). ABIDE allows assessment of whether harmonization preserves clinically meaningful group differences (controls vs. ASD) while removing scanner variance, and is also used to examine the effect of harmonization parameters (gam_df_, number of IQMs) on this preservation. Only the control subjects were used for the MAE comparison.

### Harmonization conditions compared

For each dataset, we compared brain age MAE under four harmonization conditions:

1. **Raw**: no harmonization.
2. **ComBat-GAM:** site harmonization using the GAM extension of ComBat (Pomponio et al., 2020), preserving age (and group, where applicable).
3. **ComCat (IQMs only):** harmonization with IQMs as continuous nuisance variables, no site information used.
4. **ComCat (IQMs + site):** harmonization with both IQMs and site as nuisance variables.

For ABIDE, additional analyses examined the effect of (i) the number of IQMs included as nuisance variables and (ii) the GAM basis dimension gam_df_ on harmonization performance and on the preservation of the controls-vs-ASD group difference.

For datasets without a meaningful site label (Tohoku, MR-ART), only conditions 1 and 3 apply.

## Results

### Overall harmonization performance

ComCat reduced the mean absolute error (MAE) in all five datasets, compared to the raw (unharmonized) data and ComBat-GAM (Table 5). The largest improvements were seen in the datasets where image quality variation was the dominant source of unwanted variability. In the Tohoku dataset, where all variation is acquisition-driven and no site label exists, ComCat reduced MAE from 5.001 to 0.982 years when using only IQMs, which is an 80% reduction. In the ON-Harmony dataset, MAE decreased from 4.103 (raw data) to 3.546 (ComBat-GAM) to 1.773 (ComCat with IQMs only) or 1.658 (ComCat with IQMs and site model), showing that IQMs capture substantial within-site variability that is not addressed by site harmonization alone. In the MR-ART dataset ComCat reduced MAE from 3.845 to 3.1 years when using only IQMs. The Buchert dataset exhibited the most pronounced ComBat-GAM effect (MAE=0.904) due to its robust site structure (116 scanners). However, incorporating IQMs into the site model in ComCat (IQMs + site) resulted in an additional reduction to 0.698. Figures 1-4 show the distributions of MAE for each scanner as boxplots. Figure 5 shows the distributions of MAE for the different image quality ratings as boxplots.

**Figure 1.**
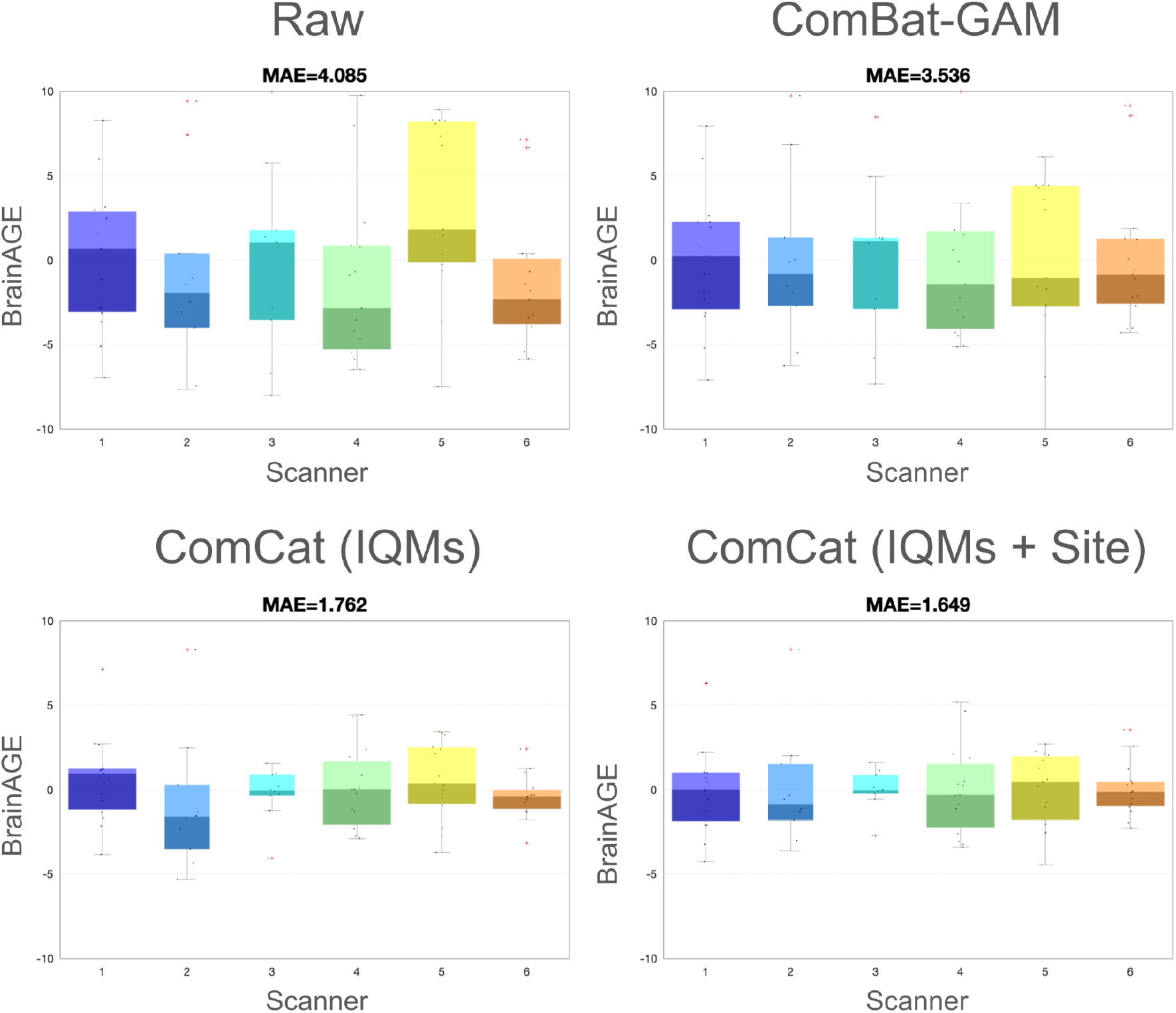
MAE in brain age prediction by harmonization condition for ON-Harmony data. The distributions of the MAE estimates for each scanner are displayed as a boxplot.

**Figure 2.**
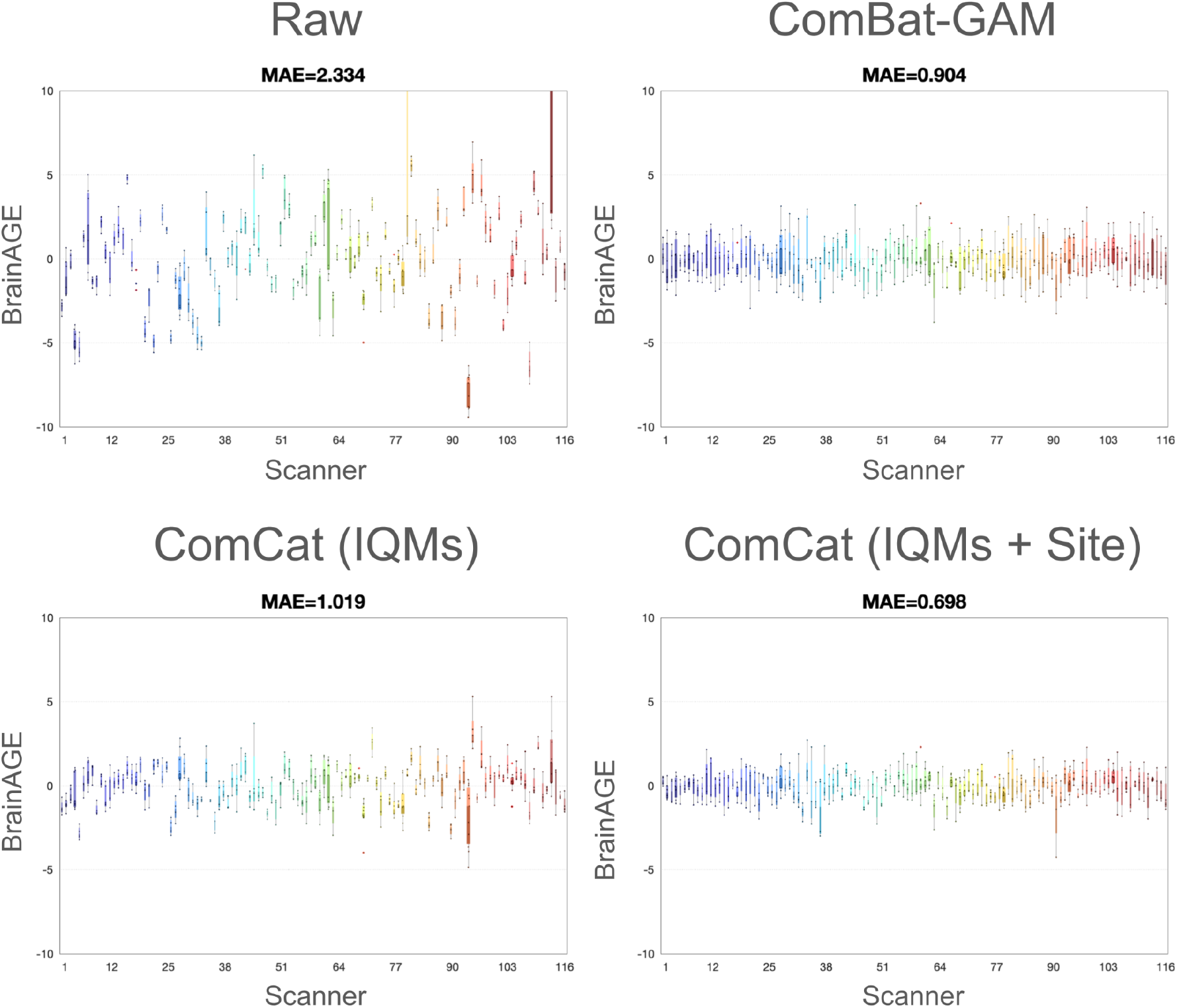
MAE in brain age prediction by harmonization condition for Buchert data. The distributions of the MAE estimates for each scanner are displayed as a boxplot.

**Figure 3.**
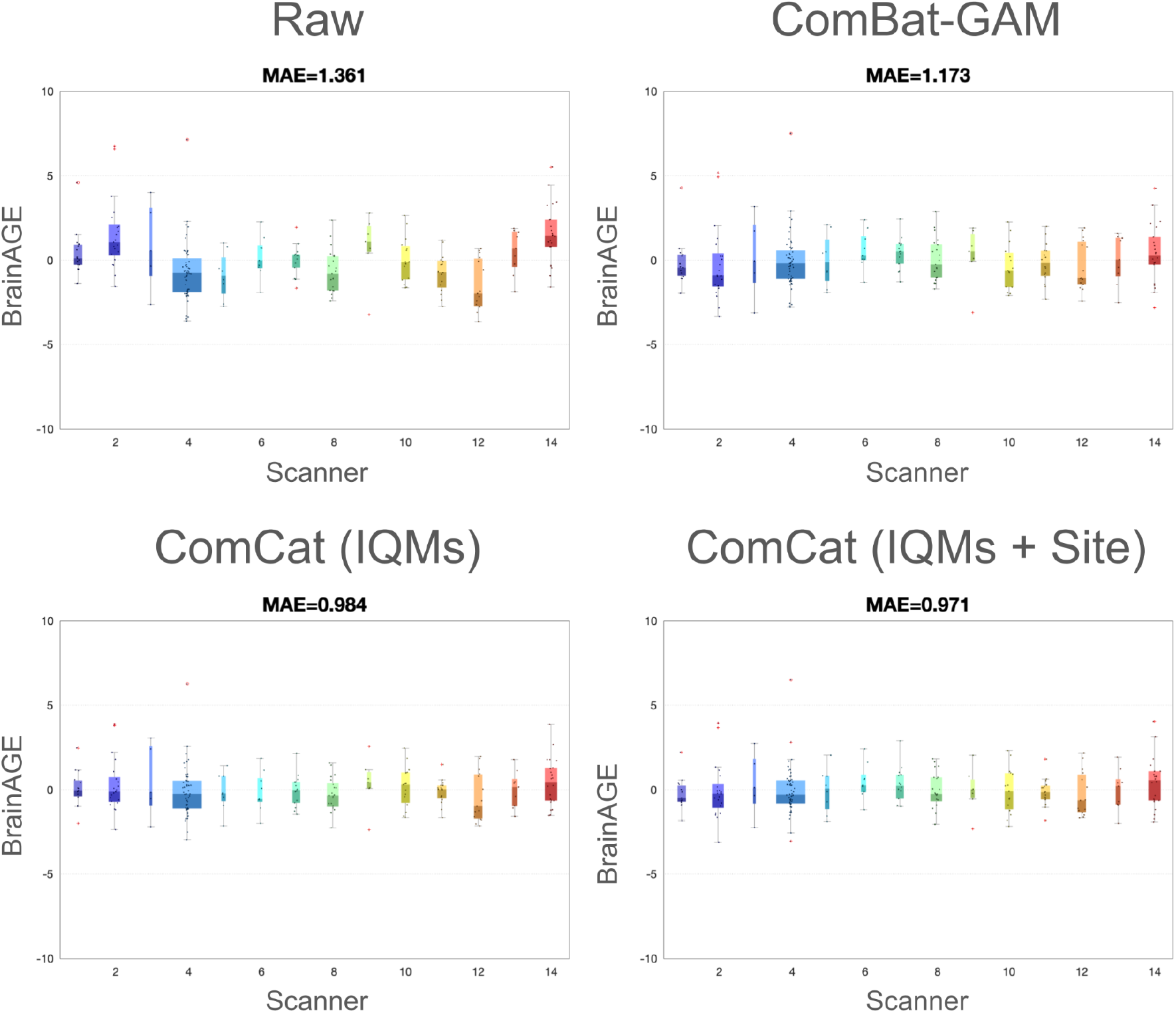
MAE in brain age prediction by harmonization condition for control subjects of ABIDE data. The distributions of the MAE estimates for each scanner are displayed as a boxplot.

**Figure 4.**
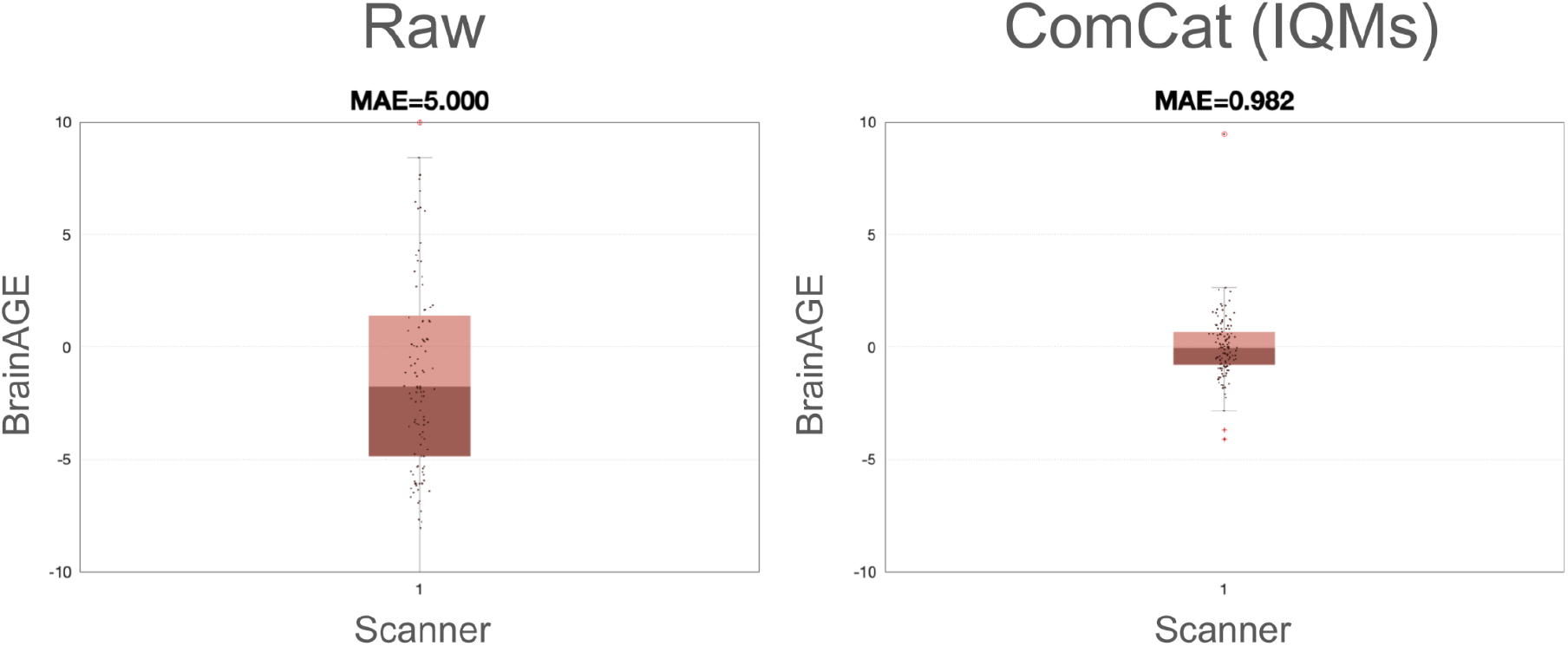
MAE in brain age prediction by harmonization condition for Tohoku data. The distributions of the MAE estimates for each scanner are displayed as a boxplot.

**Figure 5.**
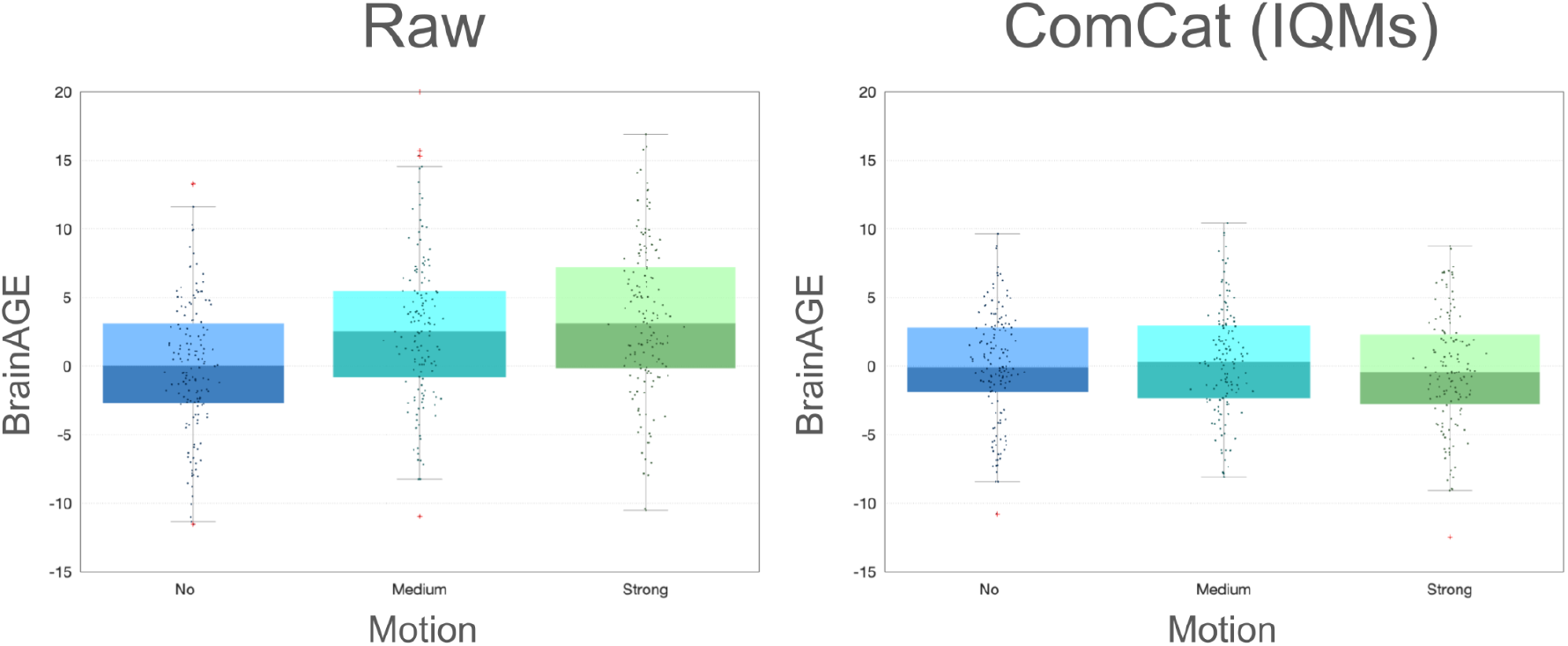
The mean predicted brain age (in years) for MR-ART scans is shown, stratified by motion level, before and after ComCat harmonization with IQMs. After harmonization, the systematic, motion-induced increase in brain age observed in the raw data is effectively eliminated.

### Effect of the number of IQMs (ABIDE)

Sequentially adding IQMs to the ComCat nuisance design progressively reduced mean absolute error (MAE) while preserving the difference between controls and ASD (Table 7). MAE of controls decreased monotonically from 1.285 (one IQM: IQR) to 0.984 (seven IQMs). The one-tailed p-value for the ASD vs. Controls test remained statistically significant from two IQMs onward. There was no consistent loss of group separation as more IQMs were added. The set of seven IQMs (IQR + SIQR + ContrastR + Res_RMS + Res_ECR + NCR + ICR) was used as the default in all subsequent analyses.

**Table 6.** The median brain age prediction (in years) by harmonization condition, differentiated by the three motion levels, is shown for the MR-ART dataset.

| Sample | Raw | ComCat (IQMs) |
| --- | --- | --- |
| No motion | -0.008 | -0.085 |
| Medium motion | 2.499 | 0.320 |
| Strong motion | 3.079 | -0.444 |

**Table 7.**
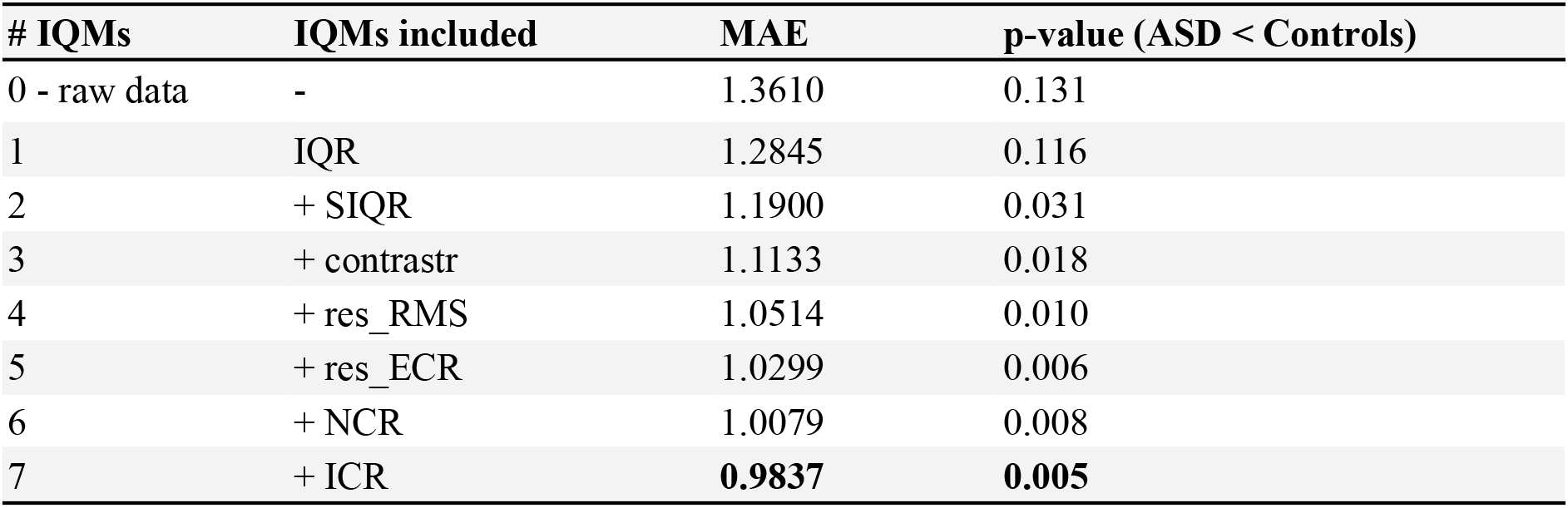
Effect of the number of IQMs on ComCat performance in ABIDE (covariates of interest: age + group).

### Effect of GAM basis dimension (ABIDE)

Increasing the B-spline basis dimension, gam_df_, reduced MAE of controls consistently but had a non-monotonic and condition-dependent effect on preserving group differences (Table 8). When both age and group were included as covariates of interest, the controls-versus-ASD p-value remained significant across all tested gam_df_ values (4 to 14). However, when only age was considered (and group was not modeled), the p-value increased from 0.018 at gam_df_ = 4 to 0.083 at gam_df_ = 14. This indicates that, with a high gam_df_, ComCat absorbed more group-related variance into the IQM smooth terms. We selected gam_df_ = 10, the upper bound of the auto-selection rule for our sample sizes, as a conservative default that captures nonlinear nuisance shape while preserving biologically meaningful group structure.

**Table 8.** Effect of GAM basis dimension gam_df_ on ComCat performance in ABIDE. The lowest value for each column is displayed in bold.

| $\text{gam}_{df}$ | Preserving<br>age + group<br>MAE | P-value<br>(ASD < Controls) | Preserving<br>age only<br>MAE | P-value<br>(ASD < Controls) |
| --- | --- | --- | --- | --- |
| 0 - ComBat-GAM | 1.1730 | 0.039 | 1.1770 | 0.041 |
| 4 | 1.0960 | 0.011 | 1.0996 | 0.018 |
| 6 | 1.0512 | 0.009 | 1.0542 | **0.021** |
| 8 | 1.0193 | 0.007 | 1.0235 | 0.022 |
| 10 | 0.9837 | **0.005** | 0.9857 | 0.022 |
| 12 | 0.9189 | 0.012 | 0.9200 | 0.046 |
| 14 | **0.8759** | 0.025 | **0.8770** | 0.083 |

## Discussion

### Main findings

ComCat improved the accuracy of predicting brain age across five datasets: controlled traveling-phantom designs, an extreme single-subject multi-scanner design, an acquisition-parameter-only single-scanner design, and a heterogeneous clinical multi-site cohort. In all scenarios, ComCat with IQMs reduced mean absolute error (MAE) relative to ComBat-GAM. Importantly, in the two designs where image quality was the dominant source of unwanted variation (Buchert: highly heterogeneous private practice scanners; Tohoku: deliberately varied acquisition parameters), ComCat outperformed ComBat-GAM, even without explicit site modeling. This demonstrates that continuous, IQM-based harmonization captures variability that discrete site labels do not. These results are consistent with recent findings that lower image quality systematically biases brain age estimates toward older ages (Nárai et al., 2024) and that scan quality and field strength affect brain measures (Gilmore et al., 2021).

The MR-ART results illustrate this most directly. In the unharmonized data, the mean predicted brain age increased systematically from -0.01 years for scans without motion, to 2.499 years for scans with medium motion, to 3.079 years for scans with strong motion. This increase mirrors the proportional motion-induced inflation reported by Moqadam et al. (2024). In their study, higher motion made participants appear neurobiologically older in direct proportion to motion severity. However, after ComCat harmonization with IQMs, this gradient was eliminated. The mean predicted brain age was 0.32 years for medium motion and -0.444 years for strong motion (Table 6). These values are near zero and are indistinguishable from the good-quality reference. Therefore, ComCat not only reduces overall MAE in heterogeneous data but also specifically removes the motion-driven systematic bias that compromises brain age estimates in motion-corrupted scans.

### MAE as a harmonization metric

We used MAE in brain age prediction as the primary evaluation metric. In test-retest and traveling-phantom designs (ON-Harmony, Buchert, and Tohoku), chronological age remains relatively consistent across scans. Therefore, all observed variation in predicted brain age predominantly reflects technical sources, such as scanner hardware, acquisition parameters, image quality, and motion artifacts. In these settings, a lower MAE is a direct measure of harmonization effectiveness because it quantifies how much technical variance has been removed. In multi-site clinical samples (ABIDE), MAE reflects the removal of scanner variance and the preservation of biological signal. A lower MAE alongside preserved group separation indicates that the harmonization is functioning properly. Thus, MAE offers a single, interpretable summary of harmonization quality sensitive to both over- and under-correction.

### GAM basis dimension and the preservation of group differences

The auto-selection of gam_df_ involves a deliberate trade-off. Higher basis dimensions produce more flexible smoothing functions of the IQMs and consequently lower MAE. In ABIDE, for example, setting gam_df_ to 14 reduced MAE to 0.876 years, compared to 0.984 with gam_df_ set to 10. However, when a group is not explicitly preserved as a covariate of interest, very flexible smooths begin to absorb biologically meaningful group-related variance into the IQM terms. With gam_df_ = 14 and only age preserved, for example, the controls-vs-ASD p-value rose to 0.083, which is no longer statistically significant. Therefore, the upper bound of gam_df_ = 10 was chosen as a conservative default that balances harmonization strength against the risk of overfitting nuisance variables and inadvertently removing biological signals. Users with samples in which group structure can be confidently included as a preserved covariate may safely increase gam_df_. Users harmonizing data without clear group labels should retain the default setting.

### Operating without site labels

ComCat’s ability to operate without site labels is valuable in practice, since scanner identity may be ambiguous (e.g., after software upgrades or coil replacements) or unavailable due to data privacy. The Buchert and Tohoku datasets show that image quality metric (IQM)-based harmonization can substantially reduce technical variance, even when conventional ComBat-style methods are not applicable. IQMs can be computed for any image, regardless of its origin, and they capture the net effect of all acquisition-related factors on the final image. This makes them a natural and broadly available basis for harmonization. In this study, we used CAT12-derived IQMs, however, the ComCat framework is agnostic to the IQM source. IQMs from MRIQC (Esteban et al., 2017) or other quality assessment pipelines could be used without modifying the algorithm.

### Relationship to other harmonization frameworks

ComCat is complementary to, not competitive with, existing extensions of the ComBat family. Each one addresses a different aspect of the harmonization problem. We compared our model with the ComBat-GAM (Pomponio et al., 2020), which extends modeling of preserved covariates (e.g., age) with smooth nonlinear terms, while keeping the nuisance model categorical. ComCat, on the other hand, extends the nuisance model by incorporating continuous confounding variables (e.g., IQMs) modeled as smooth functions. Notably, ComCat’s smooth treatment of nuisance covariates is methodologically similar to ComBat-GAM’s smooth treatment of preserved covariates, but they are applied to opposite sides of the model. Another recent model, ComBatLS (Gardner et al., 2024) extends the scale model to preserve covariate effects on variance, while CovBat (Chen et al., 2022) addresses cross-feature covariance in addition to per-feature mean and variance. These approaches modify different components of the same underlying location/scale model, and they could, in principle, be combined. Nevertheless, only a few models have been developed to consider the longitudinal data (e.g., LongComBat; Beer et al., 2020), and future work is warranted to extend ComCat’s handling of longitudinal designs.

A separate line of work has used IQMs for harmonization through a fundamentally different mechanism. Neuroharmony (Garcia-Dias et al., 2020) uses a random forest meta-model trained on a mega-dataset of 15,026 subjects to predict the volumetric corrections that ComBat would prescribe. This meta-model uses 68 MRIQC IQMs and basic demographics as input features. Although ComCat and Neuroharmony are motivated by the idea that IQMs carry information about scanner-induced variability that is otherwise inaccessible from a single image, they differ in three important respects.

First, ComCat is a direct statistical model, not a learned approximation. Location and scale corrections are derived from harmonized data via an explicit, interpretable regression on the design matrix, where all parameters can be estimated from the given sample. By contrast, Neuroharmony learns a mapping from IQMs to ComBat corrections using a training corpus, which it then applies to new data. The validity of this transfer depends on the assumption that the relationship between the IQMs and the required corrections in the new data falls within the distribution covered by the training set, a constraint that the authors explicitly acknowledge. Recent work extending Neuroharmony to clinical Alzheimer’s disease populations found that concordance dropped from over 97% on scanners seen during training to under 79% on unseen scanners (Archetti et al., 2025). This illustrates the limits of out-of-distribution generalization.

Second, ComCat models continuous nuisance variables directly within the harmonization equation. Neuroharmony, on the other hand, inherits its underlying correction structure from ComBat. Therefore, it cannot represent IQM effects beyond what ComBat applied during training. Neuroharmony’s stepwise application of ComBat (sex first, then age, then scanner) treats IQMs as only predictors of the final corrections and not sources of unwanted variation that are modeled independently. ComCat’s joint OLS estimation with smooth IQM terms enables the capture and removal of nonlinear IQM effects in a single transparent step and allows for the combination with explicit site harmonization when site labels are available. The IQMs + site condition (Table 5) shows that this combination yields further reductions in MAE beyond IQMs alone.

Third, ComCat does not require a pre-trained meta-model and is not tied to a specific feature pipeline. Currently, Neuroharmony is trained on 101 FreeSurfer regional volumes plus 68 MRIQC IQMs. Applying it to other features, such as cortical thickness, voxel-wise gray matter (GM) density, and surface metrics, or to other IQM sets, requires retraining on a comparably large corpus. ComCat, on the other hand, operates on an existing set of continuous nuisance variables. This makes ComCat directly applicable to voxel-based morphometry, surface-based morphometry, regional volumes, and brain age features without modification.

The two approaches occupy complementary niches. Neuroharmony is ideal for deploying a fixed, validated harmonization to new images from unknown scanners, particularly in clinical translational pipelines. ComCat is suited to studies where the data can be processed jointly, the IQM source or feature pipeline differs from the Neuroharmony training set, or the modeling of continuous nuisance effects needs to be transparent and tunable. Hybrid approaches, such as using ComCat-derived corrections as the basis for a Neuroharmony-style meta-model, are an interesting direction for future work.

### Scope and limitations

ComCat is presented here as a harmonization step for brain age prediction. Unlike voxel-wise statistical tests, the downstream Gaussian process regression model and cross-validated evaluation do not assume sample independence. Thus, the correlation structure introduced by two-step batch correction (Li et al., 2021) does not bias the present results. This method can also be easily applied to normative modeling frameworks that use similar machine learning evaluation paradigms. For applications in which harmonized data will be used for mass-univariate VBM or SBM testing, the correlation correction proposed by Li et al. should be combined with ComCat. This extension is straightforward in principle and will be addressed in future work.

A second limitation is that ComCat applies scale correction only for categorical batch effects. Continuous nuisance variables receive location correction, but not variance correction, as discussed in the Methods section. This is appropriate when IQMs primarily affect mean feature levels, which appears to be the dominant regime in the present datasets. However, future extensions could incorporate heteroscedasticity models for settings in which substantial IQM-dependent variance effects are present.

Despite these considerations of scope, ComCat fills a clear gap in the current harmonization landscape by providing a transparent and tunable framework for jointly correcting categorical site effects and continuous, potentially nonlinear nuisance variables. ComCat’s applicability naturally extends beyond brain age prediction to voxel- and surface-based morphometry, regional volumetric analyses, and normative modeling. It can operate with or without site labels. By explicitly modeling image quality as a continuous source of variation, ComCat provides a way to harmonize heterogeneous neuroimaging data, including data from clinical and retrospective settings where conventional, site-based methods are difficult or impossible to apply. We anticipate that this flexibility will expand the scope of multi-site studies from which reliable inferences can be drawn.

## Source Code Availability and Requirements

Project name: ComCat

Project homepage: https://github.com/ChristianGaser/ComCat

Operating system(s): Platform independent (MacOS, Linux, Windows)

Programming language: Python

Other requirements: Python (3.9-3.12) License: GPL 2.0

## References

Archetti D, et al. (2025). A Machine Learning Model to Harmonize Volumetric Brain MRI Data for Quantitative Neuroradiologic Assessment of Alzheimer Disease. Radiol Artif Intell., 7(1):e240030

Beer JC, Tustison NJ, Cook PA, Davatzikos C, Sheline YI, Shinohara RT, Linn KA (2020). Longitudinal ComBat: A method for harmonizing longitudinal multi-scanner imaging data. NeuroImage, 220, 117129.

Buchert R, et al. (2025). Frequently Traveling Human Phantom (FTHP) dataset for brain MRI harmonization. Dataset: https://www.kaggle.com/datasets/ukeppendorf/frequently-traveling-human-phantom-fthp-dat aset

Chen AA, Beer JC, Tustison NJ, Cook PA, Shinohara RT, Shou H; Alzheimer’s Disease Neuroimaging Initiative (2022). Mitigating site effects in covariance for machine learning in neuroimaging data. Human Brain Mapping, 43, 1179–1195.

Cole JH, Franke K (2017). Predicting age using neuroimaging: innovative brain ageing biomarkers. Trends in Neurosciences, 40(12), 681–690.

Dahnke R, Kalc P, Ziegler G, Grosskreutz J, Gaser C (2025). Segmentation-based quality control of structural MRI using the CAT12 toolbox. Gigascience, 14:giaf146.

Di Martino A, Yan CG, Li Q, Denio E, Castellanos FX, Alaerts K, et al. (2014). The autism brain imaging data exchange: towards a large-scale evaluation of the intrinsic brain architecture in autism. Molecular Psychiatry, 19(6), 659–667.

Esteban O, Birman D, Schaer M, Koyejo OO, Poldrack RA, Gorgolewski KJ (2017). MRIQC: Advancing the automatic prediction of image quality in MRI from unseen sites. PLOS ONE, 12(9), e0184661.

Evans AC; Brain Development Cooperative Group (2006). The NIH MRI study of normal brain development. NeuroImage, 30(1), 184–202.

Fortin JP, Parker D, Tunç B, Watanabe T, Elliott MA, Ruparel K, et al. (2017). Harmonization of multi-site diffusion tensor imaging data. NeuroImage, 161, 149–170.

Fortin JP, Cullen N, Sheline YI, Taylor WD, Aselcioglu I, Cook PA, et al. (2018). Harmonization of cortical thickness measurements across scanners and sites. NeuroImage, 167, 104–120.

Franke K, Ziegler G, Klöppel S, Gaser C; Alzheimer’s Disease Neuroimaging Initiative (2010). Estimating the age of healthy subjects from T1-weighted MRI scans using kernel methods: exploring the influence of various parameters. Neuroimage, 50(3):883–92.

Franke K, Gaser C (2019). Ten years of BrainAGE as a neuroimaging biomarker of brain aging: What insights have we gained? Frontiers in Neurology, 10, 789.

Garcia-Dias R, Scarpazza C, Baecker L, Vieira S, Pinaya WHL, Corvin A, et al. (2020). Neuroharmony: A new tool for harmonizing volumetric MRI data from unseen scanners. NeuroImage, 220, 117127.

Gardner M, Shinohara RT, Bethlehem RAI, Romero-Garcia R, Warrier V, Dorfschmidt L, et al. (2024). ComBatLS: A location- and scale-preserving method for multi-site image harmonization. Human Brain Mapping, 45, e70085.

Gaser C, Dahnke R, Thompson PM, Kurth F, Luders E; The Alzheimer’s Disease Neuroimaging Initiative (2024). CAT: a computational anatomy toolbox for the analysis of structural MRI data. GigaScience, 13, giae049.

Gilmore AD, Buser NJ, Hanson JL (2021). Variations in structural MRI quality significantly impact commonly used measures of brain anatomy. Brain Informatics, 8, 7.

Hawco C, et al. (2025). ON-Harmony: A multi-site traveling-subject dataset for harmonization benchmarking. Dataset: https://openneuro.org/datasets/ds004712.

Johnson WE, Li C, Rabinovic A (2007). Adjusting batch effects in microarray expression data using empirical Bayes methods. Biostatistics, 8(1), 118–127.

Kalc P, Dahnke R, Hoffstaedter F, Gaser C; Alzheimer’s Disease Neuroimaging Initiative (2024). BrainAGE: Revisited and reframed machine learning workflow. Human Brain Mapping, 45(3), e26632.

Kalc P, Ter Veer M, Dahnke R, Ziegler G, Kühn S, Gaser C (2026). Factors Contributing to Short-Term Structural Variability in a Longitudinal MRI Dataset. Hum Brain Mapp, 47(4):e70500.

Li T, Zhang Y, Patil P, Johnson WE (2021). Overcoming the impacts of two-step batch effect correction on gene expression estimation and inference. Biostatistics, 23(2), 635–652.

Lombardi A, Diacono D, Amoroso N, Monaco A, Tavares JMRS, Bellotti R, Tangaro S (2020). Extensive evaluation of morphological statistical harmonization for brain age prediction. Brain Sciences, 10(6), 364.

Marquand AF, Rezek I, Buitelaar J, Beckmann CF (2016). Understanding heterogeneity in clinical cohorts using normative models: Beyond case-control studies. Biological Psychiatry, 80(7), 552–561.

Moqadam R, Dadar M, Zeighami Y (2024). Investigating the impact of motion in the scanner on brain age predictions. Imaging Neurosci (Camb), 2:imag–2–00079.

Nárai, Á., Hermann, P., Auer, T. et al. (2022). Movement-related artefacts (MR-ART) dataset of matched motion-corrupted and clean structural MRI brain scans. Sci Data 9, 630.

Nárai Á, Hermann P, Auer T, Kemenczky P, Szalma J, Homolya I, et al. (2024). The effect of head motion on brain age prediction using deep convolutional neural networks. NeuroImage, 290, 120548.

Opfer R, Krüger J, Spies L, Ostwaldt AC, Kitzler HH, Schippling S, Buchert R (2023). Automatic segmentation of the thalamus using a massively trained 3D convolutional neural network: higher sensitivity for the detection of reduced thalamus volume by improved inter-scanner stability. Eur Radiol., 33(3):1852–1861

Pomponio R, Erus G, Habes M, Doshi J, Srinivasan D, Mamourian E, et al. (2020). Harmonization of large MRI datasets for the analysis of brain imaging patterns throughout the lifespan. NeuroImage, 208, 116450.

Reuter M, Tisdall MD, Qureshi A, Buckner RL, van der Kouwe AJW, Fischl B (2015). Head motion during MRI acquisition reduces gray matter volume and thickness estimates. Neuroimage, 107:107–115.

Rutherford S, Kia SM, Wolfers T, Fraza C, Zabihi M, Dinga R, et al. (2022). The normative modeling framework for computational psychiatry. Nature Protocols, 17, 1711–1734.

Smith SM, Vidaurre D, Alfaro-Almagro F, Nichols TE, & Miller K. (2019). Estimation of brain age delta from brain imaging. NeuroImage, 200, 528–539.

Thyreau B, Taki Y, Yokota S, et al. (2013). Practical impact of MRI parameters on the voxel based-morphometry measures in SPM. Presented at the 19th Annual Meeting of the Organization for Human Brain Mapping, Seattle, WA

Warrington S, Ntata A, Mougin O, et al. (2023). A resource for development and comparison of multimodal brain 3 T MRI harmonisation approaches. Imaging Neurosci (Camb), 1:imag–1–00042.

